# Anethole dithiolethione improves liver fatty acid metabolism in hamster fed high-fat diets

**DOI:** 10.1101/2020.02.18.954875

**Authors:** Chengcheng Zhao, Nannan Yu, Wenqun Li, Hualin Cai, Mouze Liu, Yanjie Hu, Yiping Liu, Mimi Tang

## Abstract

“Lipotoxicity” induced by excessive accumulation of free fatty acids (FFAs) in the liver, especially saturated FAs and their toxic metabolites, is closely related to metabolic diseases such as nonalcoholic fatty liver disease (NAFLD). Hydrogen sulfide (H_2_S), a novel gaseous signaling molecule, has been reported to have lipid-lowering effects, but its effect on FAs metabolism remains unclear. The purpose of this study was to investigate the effect and mechanisms of anethole dithiolethione (ADT, a sustained-release H_2_S donor) on hepatic FAs metabolism. ADT was administered daily for 4 weeks in male Syrian golden hamsters fed a high fat diet (HFD), and FAs profiles of liver tissues were analyzed using GC-MS. The results showed that in HFD-fed hamsters, ADT treatment significantly reduced the accumulation of saturated and monounsaturated fatty acids (C16:0, C18:0, C16:1, and C18:1n9), while increased the content of n-6 and n-3 series polyunsaturated fatty acids (C20:3n6, C20:4n6, and C22:6n3). Mechanistically, ADT obviously inhibited the overexpression of ACC1, FAS and SCD1, and up-regulated the levels of FATPs, L-FABP, CPT1α, FADS1 and FADS2. Notably, ADT evidently induced Mitofusin1 to facilitate mitochondrial fusion and optimize β-oxidation. These findings suggest that ADT plays a beneficial role by regulating the synthesis, desaturation, β-oxidation, uptake, binding/isolation, and transport of FAs. In conclusion, ADT is effective in improving liver FAs metabolic disorders and liver injuries caused by HFD.

## Introduction

Fatty acids (FAs) are indispensable sources of energy in cells, and also important bioactive mediators involved in many homeostasis processes, including metabolism and regulating inflammatory immune responses(1, 2). However, abnormal fatty acid metabolism (synthesis, desaturation, oxidation, absorption, transport) can lead to diseases such as hyperlipidemia, non-alcoholic fatty liver disease(NAFLD), diabetes, and atherosclerosis(3-6). The liver is the main metabolic organ and plays a vital role in maintaining the balance of fatty acid levels in the body. When the level of free FAs(mainly saturated palmitate) in the liver rises beyond its mitochondrial oxidation, heterotopic lipid deposition is induced, which is called “lipotoxicity”(5). There is growing evidence that lipotoxicity is mainly caused by long-chain saturated fatty acids (SFAs), such as palmitic acid (PA, C16: 0) and stearic acid (SA, C18: 0), while monounsaturated fatty acids (MUFAs) are generally less toxic and Polyunsaturated fatty acids (PUFAs) may even be protective(7-9). Three major mechanisms have been reported in palmitate and stearic acid-mediated lipotoxicity: (i) increased synthesis of harmful complex lipids such as diacylglycerol (DAG) and ceramide; (ii) impaired endoplasmic reticulum and mitochondrial function; (iii) membrane receptor-mediated inflammation, such as toll-like receptor 4 (TLR4)(5, 10-14).

Palmitic acid (C16: 0), stearic acid (C18: 0), and palmitoleic acid (C16: 1), oleic acid (C18: 1), the most common SFAs and MUFAs in the human body, are available in the diet or synthesized endogenously by the liver from carbohydrates, amino acids, and other fatty acids. The key enzymes in this process are acetyl-CoA carboxylase1 (ACC1) and FA synthase (FAS), which add seven malonyl CoAs to acetyl CoA, generating palmitic acid. Palmitic acid can be further metabolized to palmitoleic acid, stearic acid, and oleic acid, through delta-9 desaturase (stearoyl-CoA desaturase1, SCD1) and elongase. Compared with SFAs and MUFAs, animals and humans are unable to synthesize linoleic acid (LA, C18:2n6) and α-linolenic acid (ALA, C18:3n3) from precursors oleic acid(C18:1n9) due to the lack of delta-12 and delta-15 desaturases, so they must be obtained from diets. LA and ALA use the same enzyme systems (such as delta-5 and delta-6 desaturases (FADS1, FADS2), elongase) to produce PUFAs, including n-6 (arachidonic acid, AA, C20:4n6) and n-3 PUFAs (eicosapentaenoic acid (EPA, C20:5n3) and docosahexaenoic acid (DHA, C22:6n3)). N-6 and n-3 PUFAs are precursors of biologically active lipid mediator signaling molecules, including eicosanoids, which play an important role in regulating pro-inflammatory and / or anti-inflammatory / resolution processes (Figure 1).

**Figure 1.**
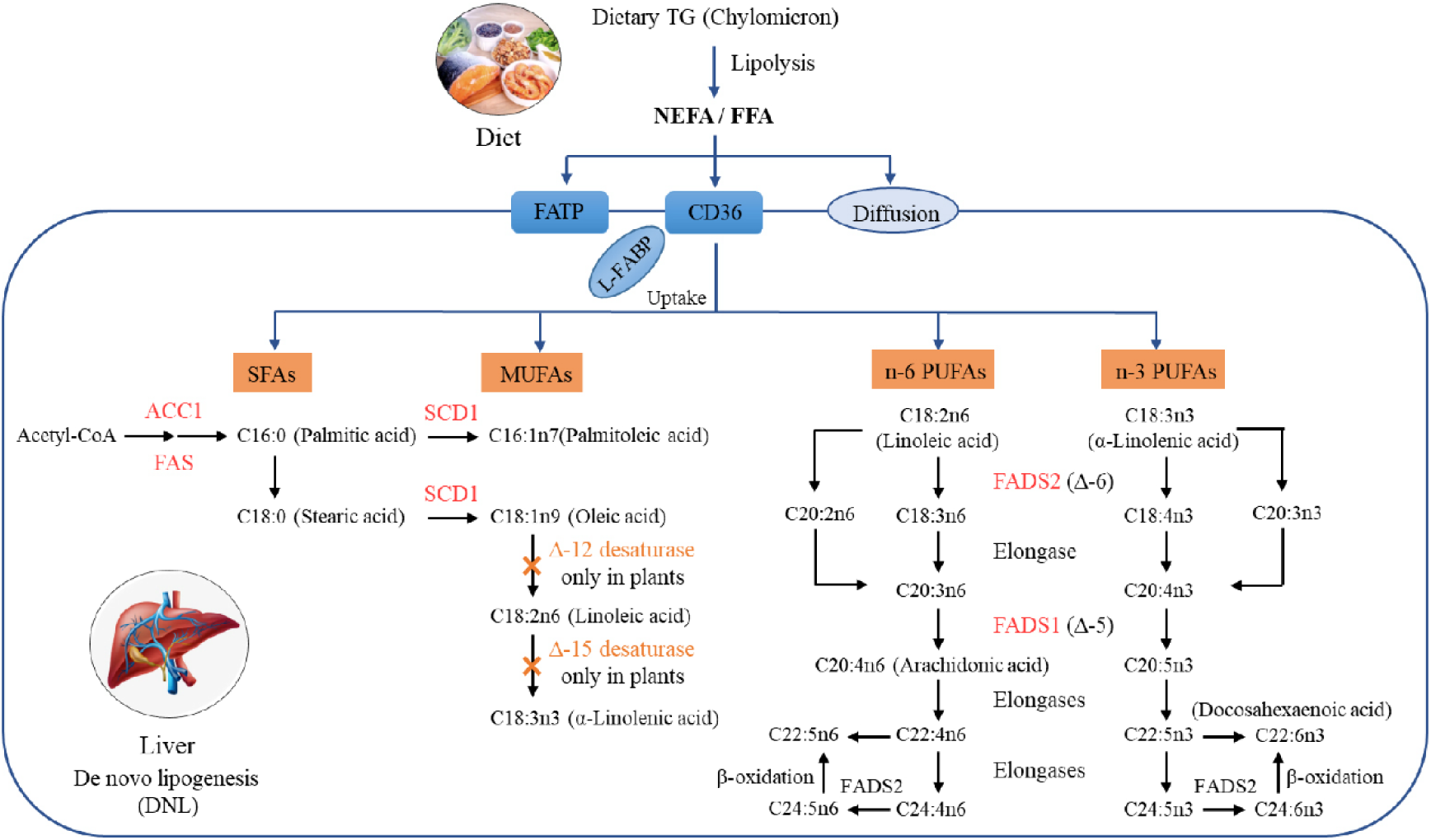
Metabolism of liver FAs. The metabolism of FAs is a complex process, involving multiple enzymes of synthesis, desaturation, elongation, and oxidation. Shown here are pathways for liver SFAs, MUFAs, n-6 and n-3 PUFAs synthesis and metabolism.

Mitochondrial β-oxidation is the most important metabolic pathway of fatty acids and is mainly regulated by rate-limiting enzymes such as carnitine palmitoyl-transferase 1α (CPT1α),which serves as a gatekeeper for fatty acids to enter mitochondria. In addition, only fused mitochondria can ensure that FAs are homogeneously distributed throughout the mitochondria, maximizing the use of FAs for β-oxidative reactions(15). However, in the mitochondrial fusion protein 1 knockout (Mitofusin1 KO) cells, mitochondria did not form a network and were fragmented, reducing the β-oxidation rate and leading to lipid accumulation in lipid droplets(15, 16).

Hydrogen sulfide (H_2_S), a well-known novel gaseous signaling molecule, is increasingly recognized as a crucial regulator of cardiovascular diseases. Studies have shown that H_2_S exerts significant protective effects on metabolic diseases (NAFLD, diabetes and atherosclerosis) through multiple properties, such as anti-inflammatory, antioxidant, inhibiting foam cell formation, improving endothelial function, and increasing insulin receptor sensitivity(17-20). H_2_S has been reported to have a lipid-lowering effect by activating liver autophagy(21), but its effect on fatty acid metabolism remains unclear. NaHS, a so-called immediate-release H_2_S donor, releases excessive H_2_S instantaneously and therefore does not mimic the production of endogenous H_2_S(22). Anethole dithiolethione (ADT; 5-(4-methoxyphenyl)-3H-1,2-dithiole-3-thione), clinically used as a hepatoprotective and choleretic drug, is also a prodrug of H_2_S and releases H_2_S in a controlled manner within a few hours in the body(23, 24). This article investigates how ADT affects hepatic fatty acid metabolism and explores its possible mechanisms.

## Materials and Methods

### 2.1 Drug

ADT(Lot.N1017A) was purchased from Dalian Meilun Biotechnology Co., LTD., and its clinical dosage is 75mg per day. The dosage of ADT in Syrian golden hamsters was calculated by human equivalent dose (HED) based on body surface area(25), with a conversion dose of 9.25mg/kg. Therefore, the dose gradient of ADT in this study was set as: 5mg/kg, 10mg/kg, 20mg/kg and 40mg/kg. ADT is insoluble in water, suspended in 0.5% sodium carboxymethylcellulose (CMC-Na) and 0.5% soybean lecithin, administered daily by gavage.

### 2.2 Animals and Diets

This study was reviewed and approved by the Laboratory Animal Care and Welfare Committee of Central South University(Approval No. 2018sydw0215). Fifty-four male Syrian golden hamsters(80-100g) were purchased from Beijing Vital River Laboratory Animal Technology Co., Ltd. (Qualified Certificate No. SCXK Jing 2016-0011). Hamsters were housed in IVC cages and free access to diet and water, which subjected to a 12-hour light/dark cycle with a relative humidity of 50% ±10% and a temperature of 22°C-25°C.

Trophic Animal Feed High-Tech Co. Ltd provided high-fat diets and control diets, both of which are formulated based on purified ingredients. The control diet was designed to meet all of the hamster’s nutritional requirements. The cholesterol and lard in the high-fat diet were 2/1000 g and 117/1000 g, respectively.

### 2.3 Experimental design

After adaptation for one week, 54 hamsters were randomly divided into six groups, with 9 animals in each group. (1) control group: control diet with vehicle (0.5% CMC-Na and 0.5% soybean lecithin) treatment; (2) HFD group: high-fat diet with vehicle (0.5% CMC-Na and 0.5% soybean lecithin) treatment; (3-6) HFD+ADT group: high-fat diet with different doses of ADT. ADT and/or vehicle were administered to the hamster by gavage once a day for 4 weeks. After 4 weeks, all hamsters fasted overnight and were euthanatized at the end of the experiment. Blood samples were collected from the heart and kept at room temperature for 1h before centrifugation at 3500 rpm for 10min. The serum was separated and stored at - 80°C until analysis. The left lobe of liver was fixed in 4% paraformaldehyde solution, and the rest was stored at - 80°C after quick-freezing by liquid nitrogen.

### 2.4 Biochemical and Histological Analysis

Serum glutamate alanine aminotransferase (ALT), aspartate aminotransferase (AST) and total bile acid (TBA) were measured by automatic biochemical analyzer(Hitachi 7600-210) in the laboratory of Second Xiangya Hospital. The fixed liver tissue was embedded in paraffin blocks, and 10-μm-thick slices were cut and stained with hematoxylin and eosin (HE) to observe histopathological changes.

### 2.5 Sample pretreatment and FFAs determination

FFAs in liver tissue were extracted and measured by GC-MS (Agilent 7890 A / 5975 C) according to the method described by Tang et al(26). In short, 750μl of dichloromethane and methanol (CH_2_Cl_2_ : CH_3_OH=1:2) mixture was added to liver tissue and homogenized, followed by centrifugation to obtain the supernatant. Then, 100μM butylated hydroxytoluene (BHT, to prevent lipid peroxidation), 250μl CH_2_Cl_2_ and 250μl water were sequentially added to the supernatant and vortexed for 30s. We transferred the lower phase, centrifuged again to take the supernatant which then evaporated to dryness under nitrogen. The residue was dissolved in n-hexane, 10μl internal standard (heptadecanoic acid) and 2ml 0.5M KOH-MeOH were added, followed by water bath heating at 60°C for 20min. After cooling for 10min, 3ml of 12.5% H_2_SO_4_ in methanol solution (to methylate the sample) was added and heated again at 60°C in the water bath for 1h. After cooling the sample vial, 2ml n-hexane and 1ml saturated sodium chloride (NaCl) solution were added, and let stand for 10min to stratify. The supernatant (hexane fraction) was then transferred for GC-MS analysis.

GC-MS analysis was performed on Agilent 7890 A/5975 C, with specific parameter settings modified according to the previously reported procedures. The samples were separated by VF-23 ms chromatographic column (Agilent: 30m(length), 0.25mm(inner diameter), 0.25μm(film thickness)). FFA composition was determined based on the retention time of validated fatty acid methyl ester standard (Supelco 37, sigma). FFA content was expressed as a percentage of peak area.

### 2.6 RNA extraction, reverse transcription and real-time quantitative PCR (RT-qPCR)

The expression of 12 genes involved in liver fatty acid metabolism (synthesis, desaturation, uptake, transport and oxidation) was determined by RT-qPCR. Total RNA was extracted from frozen liver tissue with TRIzol reagent (Invitrogen) and reverse-transcribed using PrimeScript™ RT reagent kit (Takara BIO Inc., Code No.RR047A) according to manufacturer’s instructions.

RT-qPCR was performed using TB Green® Premix Ex Taq ™ II (Tli RNaseH Plus) (Takara BIO Inc., Code No. RR820A) and LightCycler® 96 system. The total volume of RT-qPCR reaction was 10µL, and its cycling parameters were set as follows: pre-denaturation for 30s at 95°C, followed by 40 cycles at 95°C for 5s and 60°C for 60s, then annealing and elongation. The expression level of each gene relative to GAPDH was determined by the 2-ΔΔCt method, and then normalized to the corresponding control group. Gene-specific primers of hamsters for RT-qPCR are listed in table 1.

**Table 1.**
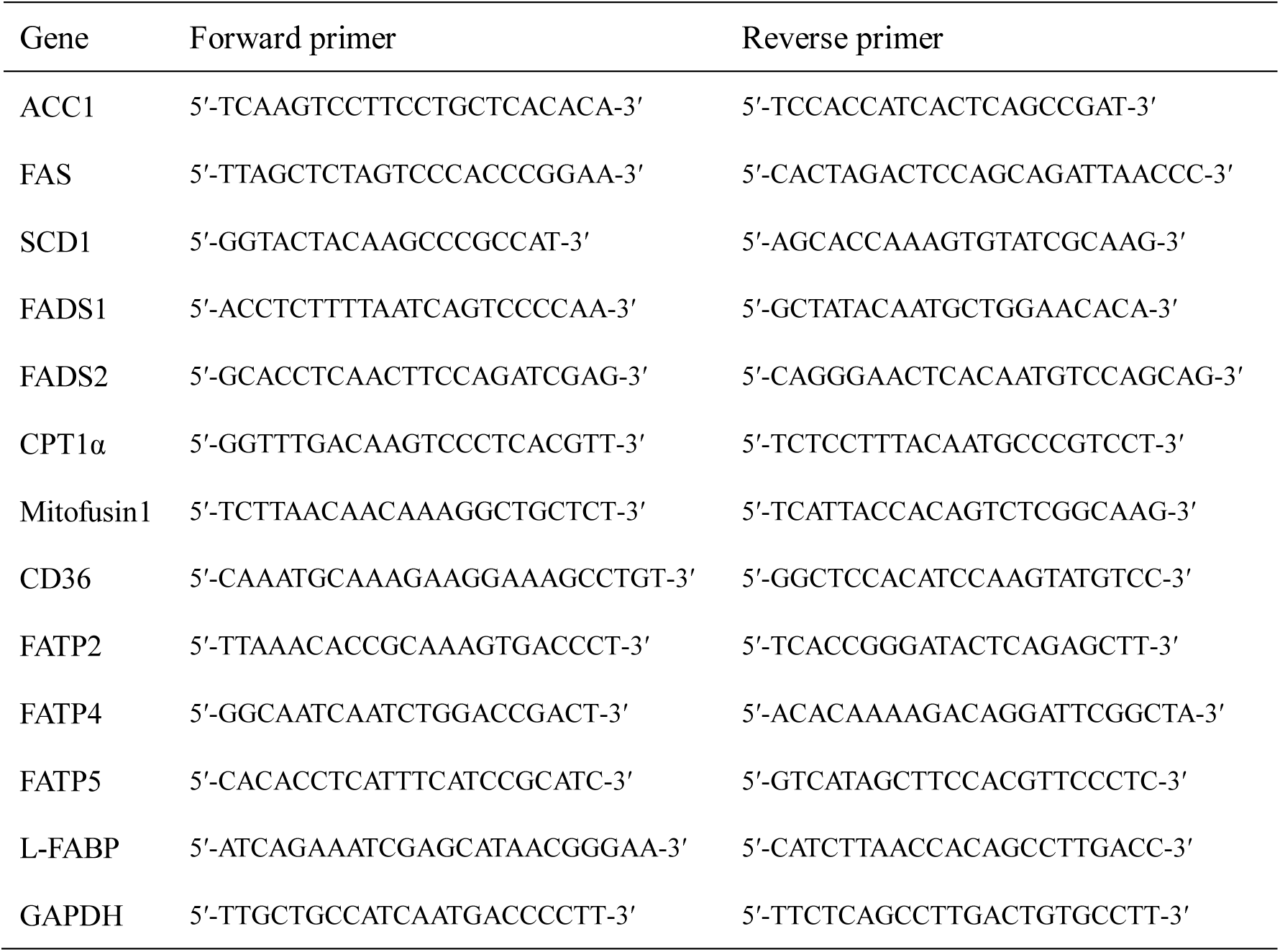
Hamster primer sequences used in this study.

### 2.7 Statistical Analysis

Statistical comparisons of the data were performed by SPSS 18.0 software using one-way analysis of variance(ANOVA). The experimental results were presented as mean ± standard deviation (SD). P values <0.05 were considered statistically significant.

## Results

### 3.1 Effect of ADT on biochemical parameters and histology

Serum ALT, AST and TBA levels were measured to assess the effect of ADT on liver function. As shown in Figure 2A, different doses of ADT significantly reduced elevated ALT, AST, and TBA levels in hamsters fed a high-fat diet. In addition, changes in these biochemical indicators have been confirmed by pathological changes in the liver. H&E staining results revealed that significant swelling of hepatocytes and increased lipid droplets in the liver of HFD-fed hamsters, while control hamsters displayed normal liver histology. Compared with the HFD group, liver steatosis in ADT intervention groups with different concentrations were improved to varying degrees, and liver structure tended to normal (Figure 2B).

**Figure 2.**
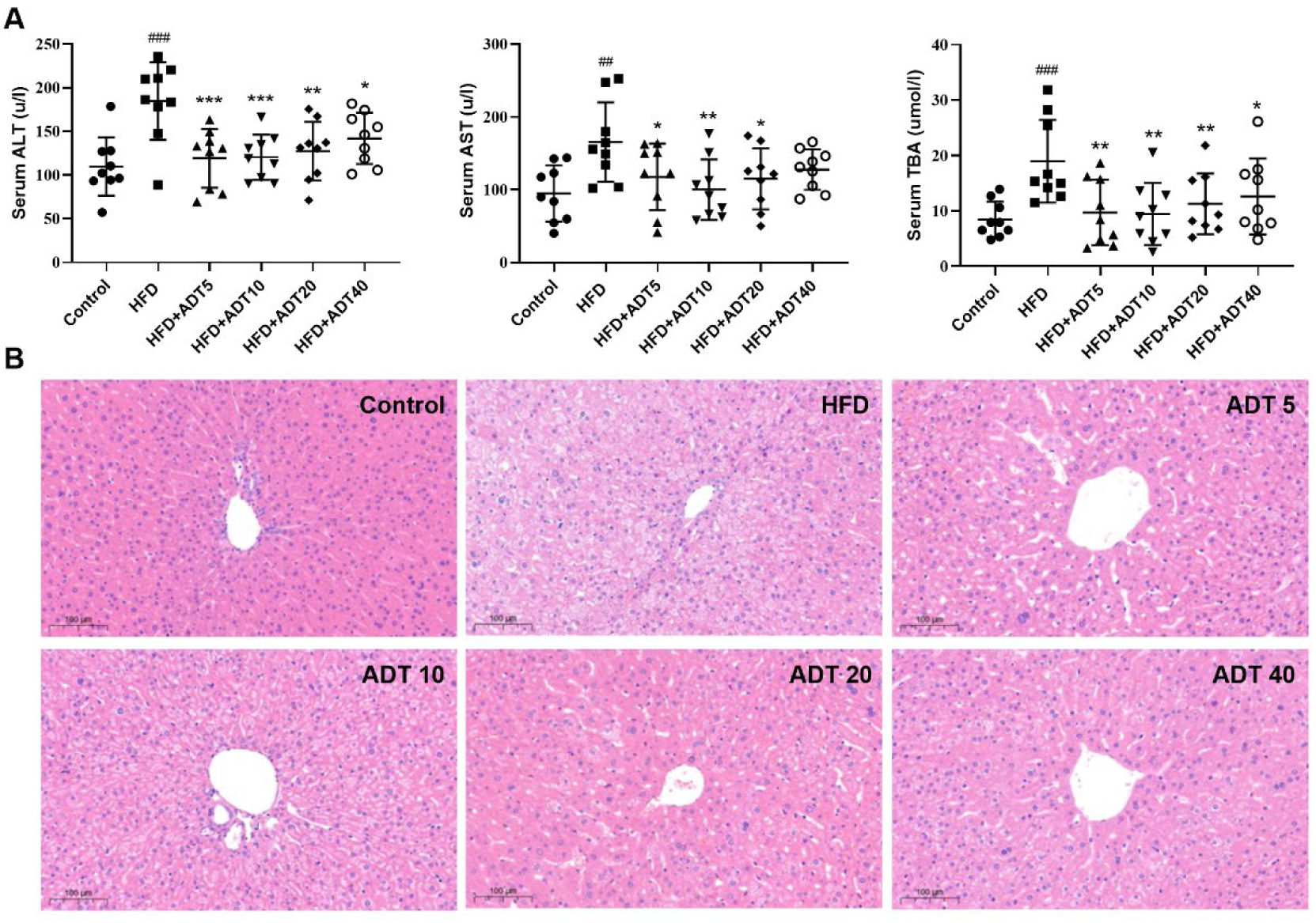
Effect of ADT on liver function of HFD-fed hamsters. (A): Serum levels of ALT, AST, and TBA in each group of hamsters (n=9). Data are presented as mean ± *SD*. ALT = alanine aminotransferase; AST = aspartate aminotransferase; TBA = total biliary acid. ^#^*p* < 0.05, ^##^*p* < 0.01, and ^###^*p* < 0.001 versus control. ^*^*p* < 0.05, ^**^*p* < 0.01, and ^***^*p* < 0.001 versus HFD. (B): Histopathological examination of hamster liver sections by HE staining (magnification, ×100).

### 3.2 Quantitative changes of FFAs in hamster liver treated with ADT

Quantitative changes of 13 FFAs detected by GC-MS in hamster liver in different experimental groups are shown in table 2 (n =9). These data indicated that total SFA and MUFA increased significantly in the HFD group compared to the control group. However, there was no obvious change in total PUFA between the two groups, with a slight increase in n-6 fatty acids and a decrease in n-3 fatty acids. ADT treatment significantly reduced total SFA and MUFA in the liver of hamsters fed a high-fat diet, and evidently up-regulated n-3 PUFA.

**Table 2.** Quantitative changes of liver FFAs in hamsters treated with ADT (*n* = 9)

| Fatty acids<br>(mg/ml) | Control | Model | ADT5<br>5mg/kg | ADT10<br>10mg/kg | ADT20<br>20mg/kg | ADT40<br>40mg/kg |
| --- | --- | --- | --- | --- | --- | --- |
| C14: 0 (myristic acid) | 0.07±0.02 | 0.09±0.05 | 0.07±0.02 | 0.07±0.01 | 0.07±0.02 | 0.06±0.01 |
| C16:0 (PA, palmitic acid) | 6.71±1.27 | 8.82±1.71<sup>###</sup> | 6.81±1.08<sup>***</sup> | 5.80±0.70<sup>***</sup> | 7.03±0.67<sup>**</sup> | 7.04±0.70<sup>**</sup> |
| C18:0 (SA, stearic acid) | 4.76±0.87 | 5.77±0.92<sup>##</sup> | 5.66±0.39 | 4.72±0.54<sup>**</sup> | 5.65±0.54 | 6.06±0.76 |
| Total SFA | 11.54±1.81 | 14.68±2.32<sup>###</sup> | 12.54±1.44<sup>**</sup> | 10.59±1.21<sup>***</sup> | 12.76±1.09<sup>*</sup> | 13.16±1.20<sup>*</sup> |
| C16:1 (palmitoleic acid) | 0.38±0.18 | 0.89±0.23<sup>###</sup> | 0.69±0.17<sup>*</sup> | 0.59±0.15<sup>***</sup> | 0.59±0.12<sup>***</sup> | 0.52±0.07<sup>***</sup> |
| C18:1n9 (oleic acid) | 5.30±1.36 | 14.78±2.67<sup>###</sup> | 10.71±1.44<sup>***</sup> | 9.32±1.42<sup>***</sup> | 9.89±1.31<sup>***</sup> | 10.03±1.28<sup>***</sup> |
| C20:1 (eicosenoic acid) | 0.03±0.01 | 0.08±0.03<sup>###</sup> | 0.06±0.01<sup>*</sup> | 0.06±0.01<sup>*</sup> | 0.06±0.01<sup>*</sup> | 0.07±0.02 |
| C24:1 (nervonic acid) | 0.09±0.03 | 0.08±0.02 | 0.11±0.05 | 0.10±0.01 | 0.11±0.05 | 0.14±0.05 |
| Total MUFA | 5.80±1.53 | 15.83±2.90<sup>###</sup> | 11.57±1.60<sup>***</sup> | 10.07±1.56<sup>***</sup> | 10.65±1.41<sup>***</sup> | 10.76±1.32<sup>***</sup> |
| C18:2n6 (LA, linolenic acid) | 5.89±1.88 | 7.43±1.34<sup>##</sup> | 6.73±1.12 | 5.64±0.79<sup>**</sup> | 6.69±0.88 | 6.88±0.59 |
| C20: 2n6 (eicosadienoic acid) | 0.04±0.01 | 0.06±0.01<sup>##</sup> | 0.06±0.01 | 0.06±0.01 | 0.06±0.01 | 0.08±0.02<sup>*</sup> |
| C20: 3n6 (DGLA, dihomogamma<br>linolenic acid) | 0.22±0.04 | 0.17±0.06 | 0.23±0.02 | 0.21±0.03 | 0.26±0.05 | 0.29±0.05<sup>**</sup> |
| C20:4n6 (AA, arachidonic acid) | 2.41±0.40 | 2.35±0.37 | 2.72±0.35<sup>*</sup> | 2.56±0.43 | 2.79±0.30<sup>*</sup> | 2.94±0.39<sup>**</sup> |
| Total n-6 PUFA | 8.57±2.24 | 10.03±1.22<sup>#</sup> | 9.74±1.41 | 8.47±0.85<sup>*</sup> | 9.81±1.12 | 10.19±0.84 |
| C18:3n3 (ALA, α-linolenic acid) | 0.07±0.05 | 0.10±0.07 | 0.07±0.03 | 0.06±0.02 | 0.06±0.02 | 0.05±0.01 |
| C22:6n3 (DHA, docosahexaenoic<br>acid) | 1.49±0.30 | 1.14±0.15<sup>##</sup> | 1.42±0.19<sup>**</sup> | 1.37±0.23<sup>*</sup> | 1.43±0.19<sup>**</sup> | 1.57±0.23<sup>***</sup> |
| Total n-3 PUFA | 1.55±0.32 | 1.23±0.13<sup>##</sup> | 1.49±0.21<sup>*</sup> | 1.43±0.23 | 1.49±0.19<sup>*</sup> | 1.61±0.22<sup>**</sup> |
| Total PUFA | 10.12±2.45 | 11.26±1.18 | 11.23±1.59 | 9.90±0.94 | 11.30±1.26 | 11.80±0.97 |
| C16:1/C16:0 | 0.06±0.02 | 0.10±0.03<sup>###</sup> | 0.10±0.03 | 0.10±0.02 | 0.08±0.02 | 0.07±0.01<sup>**</sup> |
| C18:1/C18:0 | 1.15±0.36 | 2.69±0.44<sup>###</sup> | 1.90±0.27<sup>***</sup> | 1.98±0.29<sup>***</sup> | 1.76±0.29<sup>***</sup> | 1.68±0.30<sup>***</sup> |
| C20:4n6/C18:2n6 | 0.44±0.11 | 0.33±0.08<sup>##</sup> | 0.41±0.04<sup>*</sup> | 0.46±0.12<sup>**</sup> | 0.42±0.04<sup>*</sup> | 0.43±0.06<sup>*</sup> |
| C22:6n3/C18:3n3 | 34.23±15.37 | 17.84±10.76<sup>##</sup> | 23.32±9.48 | 25.88±8.04 | 27.78±9.49 | 32.67±10.8<sup>**</sup> |

Results are presented as mean ± *SD*. SFA correspond to 14:0, 16:0 and 18:0. MUFA
correspond to 16:1, 18:1n9, 20:1 and 24:1. n-6 PUFA include 18:2n6, 20:2n6, 20:3n6 and
20:4n6; n-3 PUFA include 18:3n3 and 22:6n3; PUFA correspond to n-6 and n-3 PUFA.
<sup>#</sup> $p < 0.05$ , <sup>##</sup> $p < 0.01$ , and <sup>###</sup> $p < 0.001$ versus control. <sup>\*</sup> $p < 0.05$ , <sup>\*\*</sup> $p < 0.01$ , and <sup>\*\*\*</sup> $p < 0.001$
versus HFD.

The specific changes of different FFAs between the experimental groups were further analyzed. C16:0 as well as C18:0, the two main SFAs, were increased in HFD, and this increase was subsequently inhibited by ADT treatment. C18:1n9, the main MUFA in the body, was significantly elevated in HFD, with concentrations nearly tripling compared to control levels, which can be reduced by ADT administration. For these three FFAs, ADT at 10mg/kg was the most effective. In addition, C16:1 was also obviously up-regulated in HFD and was reduced in a dose-dependent manner by ADT. Meanwhile, the essential fatty acids (EFAs) including C18:2n6 and C18:3n3 were increased in HFD group and decreased in HFD + ADT group. In contrast, HFD-fed hamster livers showed lower PUFAs levels, including C20:3n6, C20:4n6 and C22:6n3, which were markedly elevated after ADT administration. Other FFAs (C14:0, C20:1, C20:2, and C24:1) had very low liver concentrations, C14:0 and C24:1 did not differ among the groups, while C20:1 and C20:2 were evidently altered.

### 3.3 Effect of ADT on mRNA expression of fatty acid metabolism gene in liver

To investigate the molecular mechanism of ADT, we further quantitatively measured mRNA expression levels of genes related to FA synthesis, desaturation, β-oxidation, uptake and transport in the liver (Figure 3). The expression of ACC1 and FAS was significantly increased in the HFD group and decreased dose-dependently in HFD + ADT groups (Figure 3A).

**Figure 3.**
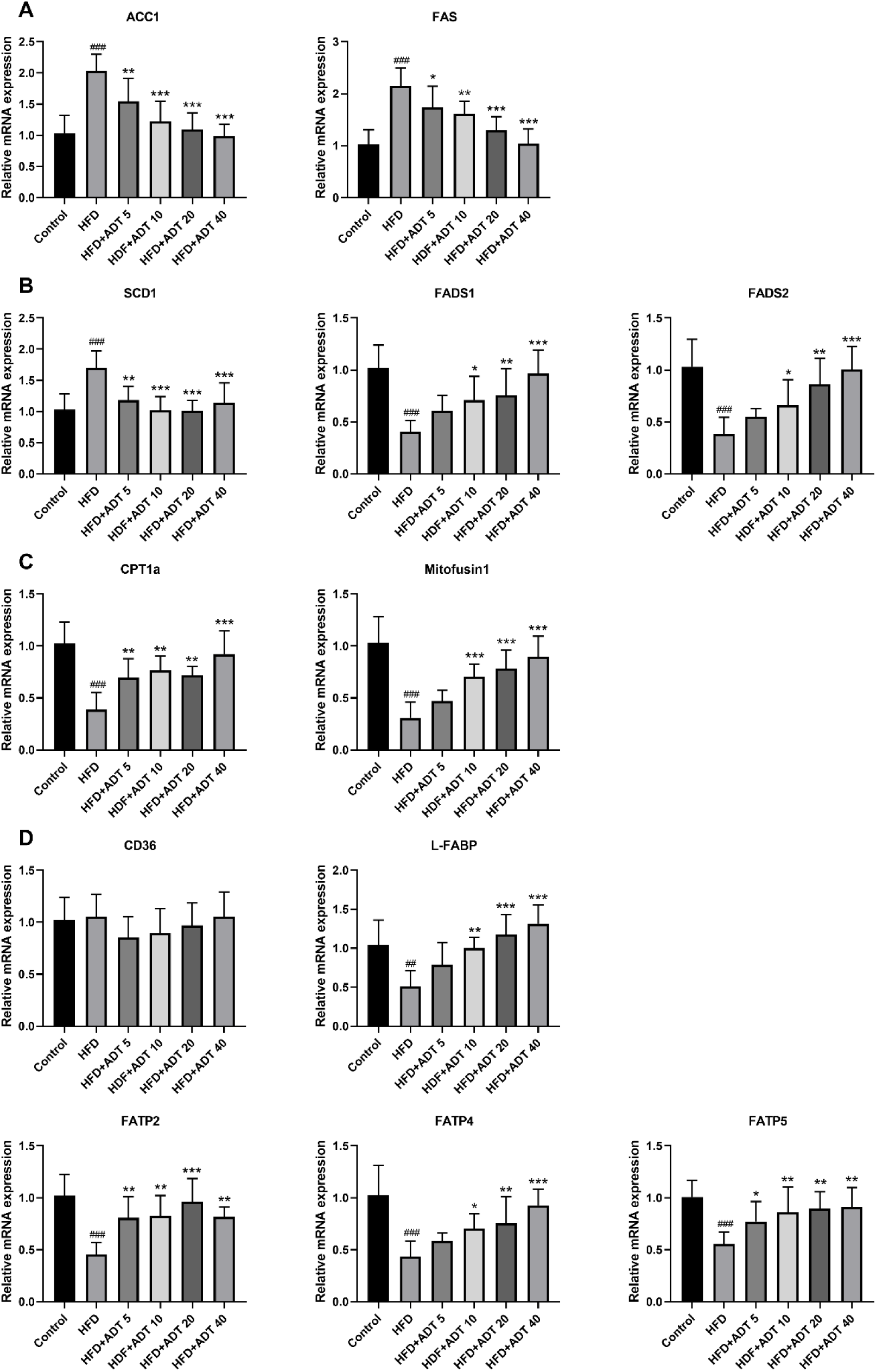
mRNA levels of genes related to FAs metabolism in hamster liver (n=6). (A). Fas synthesis: ACCl and FAS. (B). FAs desaturation: SCD1 converts SFA into MUFA, with the main product is C18:1n9; FADS1 and FADS2 metabolize EFAs(LA and ALA) into more unsaturated long-chain PUFAs. (C). FAs β-oxidation: CPT1α catalyzes FAs into mitochondria; Mitofusin1 regulates the fusion state of mitochondria. (D). FAs uptake and transport: CD36, FATP2, FATP4, FATP5 and L-FABP. ^#^*p* < 0.05, ^##^*p* < 0.01, and ^###^*p* < 0.001 versus control. ^*^*p* < 0.05, ^**^*p* < 0.01, and ^***^*p* < 0.001 versus HFD.

SCD1, a key rate-limiting enzyme that desaturates SFA into MUFA, was significantly elevated in the HFD group and attenuated by ADT administration (Figure 3B). The changes of SCD1 were associated with changes in C16:1/C16:0 and C18:1n9/C18:0 ratios (Table. 2). On the contrary, FADS1 and FADS2, key rate-limiting enzymes in the desaturation of PUFA, were obviously down-regulated in HFD and dose-dependently increased by ADT treatment (Fig. 3B). Similarly, the changes in FADS1 and FADS2 were consistent with changes in C20:4n6/C18:2n6 and C22:6n3/C18:3n3 ratio(Table. 2).

CPT1α and Mitofusin1, the regulators of fatty acid β-oxidation, were concurrently decreased in HFD group, but significantly raised in the HFD + ADT group (Figure 3C).

Moreover, genes involved in liver FA uptake and transport showed that the expression of CD36 was not obviously different among the groups, while the expression of FATP2, FATP4, FATP5 and L-FABP was significantly reduced in HFD group but activated by ADT therapy (Figure 3D).

## Discussion

In this study, we investigated the effects and mechanisms of ADT on FA metabolism. The results showed that ADT administration significantly reduced the concentrations of SFA and MUFA in HFD hamster liver, and increased the levels of n-3 PUFA. Further mechanism studies indicated that ADT treatment reduced FA synthesis, SFA desaturation, increased PUFA desaturation and promoted FAs absorption and β-oxidation, which may improving FA metabolism.

Excessive accumulation of free SFAs (Palmitic acid, PA, C16:0 and Stearic acid, SA, C18:0) in the liver can induce lipotoxicity, leading to cellular dysfunction and death(7). In our research, HFD-fed hamsters showed overexpression of ACC1 and FAS, meaning increased endogenous synthesis of PA and SA. When the increased intracellular PA level exceeds its β-oxidation in mitochondria, it is converted to harmful complex lipids such as diacylglycerin (DAG) and ceramide(5, 27). These harmful FA-derived intermediates and excessive PA provoke increased reactive oxygen species (ROS) production, damage the function of mitochondria-related ER membrane (MAM), and induce mitochondrial fragmentation, resulting in mitochondrial dysfunction and loss of ATP production(28-31). Recently, Rambold et al. reported that fragmented mitochondria failed to efficiently take up FA, causing a reduction in β-oxidation rates and further exacerbating the accumulation of FA in Mitofusin1 KO cells(15). Notably, this highly fused mitochondrial requirement was specific to β-oxidation, as opposed to glutamine oxidation, which was not affected by mitochondrial morphology(15). Our results indicated that ADT administration significantly inhibited ACC1 and FAS, up-regulated CPT1α and Mitofusin1, thereby effectively reduced the accumulation of PA and SA in HFD-fed hamsters.

Oleic acid(OA, C18: 1n9), an important MUFA produced by SCD1 metabolizing SFA(32), was significantly elevated in HFD hamsters, which may be due to the body’s adaptive results in the face of on-going metabolic stress. To counteract SFA-induced lipotoxicity, cells usually employ two main methods: 1) increasing the β-oxidation of FAs in mitochondria; and 2) inducing the storage of FAs in neutral LD (lipid droplets)(33-35). However, as mentioned above, mitochondrial β-oxidation rate decreased in HFD hamsters. The OA is mainly incorporated into relatively inert triacylglycerol (TAG) and stored in lipid droplets(5, 36), so metabolizing SFA to OA via SCD1 reduces the excessive formation of toxic lipid intermediates (DAG and ceramides)(37, 38). Nevertheless, this relative protection by channeling SFA into less toxic lipid pool is only temporary, as OA is less lipotoxic but more steatotic than PA(8, 39). Continuous overproduction of OA results in liver lipid accumulation, steatosis, and low-grade chronic inflammation, thereby potentiating the metabolic syndrome(8, 39). In our study, ADT treatment in HFD hamsters not only effectively reduced SFAs but also significantly decreased OA.

Essential fatty acids (EFA), LA (C18:2n6) and ALA (C18:3n3) are converted to their respective PUFA metabolites by the action of FADS1 and FADS2(41). LA is converted to DGLA (C20:3n6) and AA (C20:4n6), while ALA is converted to form EPA (C20:5n3) and DHA (C22:6n3). Excessive intake of HFD reduces the levels of AA, EPA and DHA(42), enhances the production of pro-inflammatory arachidic acids and ROS, leading to pro-inflammatory status(42, 43). Our results also confirmed that HFD impaired the desaturated metabolic pathway of EFAs: the ratio of AA/LA and DHA/ALA in HFD hamsters was down-regulated, and the expressions of FADS1 and FADS2 were significantly decreased. In this study, the liver concentration of EPA was too low for accurate quantification. ADT administration obviously up-regulated FADS1 and FADS2, increasing the conversion of EFA to AA and DHA.

In addition to FA synthesis, β-oxidation and desaturation, the effective uptake and channeling of exogenous FA are critical to the level of FFA in hepatocytes. Evidence is emerging that specific protein transport systems are the main mediators of transmembrane FAs-trafficking into hepatocytes, including fatty acid transposase (FAT/CD 36), fatty acid transport proteins (FATPs) and liver fatty acid binding protein (L-FABP)(44). In this study, there was no obvious difference in CD36 gene expression among the groups, while the expression of FATP2, FATP4, FATP5, and L-FABP was significantly decreased in HFD group but elevated by ADT treatment. FATP2 and FATP5 are the two main FATPs in the liver(45, 46). Studies have shown that sustained protein-mediated liver LCFAs uptake contributes to NFALD in cases of lipid oversupply(47). Liver specific FATP2 or FATP5 knockdown based on adeno-associated virus (AAV) significantly reduces hepatic steatosis induced by continuous high-fat feeding, thereby improving NAFLD(48, 49). However, it is worth noting that recent clinical studies have reported that low expression of FATP5 is the most significant risk factor for liver fat loss(50) and is linked to the progression of NAFLD(50, 51). Different from FATP2 and FATP5, FATP4 is localized in the liver endoplasmic reticulum and drives FAs uptake indirectly by vectorial acylation rather than acting as a transporter itself(52). Adipose or hepatocyte-specific FATP4 deficient mice under high fat/sugar diets exhibited a higher degree of hepatic steatosis(53, 54).

Whether increased FATP-mediated LCFAs uptake is salutary or detrimental may depend on duration, tissue type, and subsequent fate of FAs. Upon entering hepatocytes, LCFAs and LCFA-CoA are bound/isolated by L-FABP to minimize the toxic effects of excess free FAs(55). L-FABP is a vital endogenous cytoprotectant that transports bound LCFAs for rapid removal in various cellular compartments(56-59), including endoplasmic reticulum, lipid vesicles, peroxisomes, mitochondria, and nucleus. More notably, L-FABP is not only the intracellular counterpart of albumin but also has strong antioxidant properties(56, 60, 61). Studies have shown that the loss of L-FABP may enhance the production of lipotoxic inflammatory mediators, impair the oxidative pathway of FAs, and render hepatocytes more vulnerable to the harmful effects of LCFAs, thus promoting the development of NAFLD(62, 63). Consistent with these studies, the expression of L-FABP and CPT1α was significantly reduced in HFD-fed hamsters, which suggested that delivery of FAs to mitochondria was inhibited, thereby impairing oxidation and leading to overaccumulation of FAs. ADT treatment significantly up-regulated the expression of L-FABP and CPT1α, therefore improving the β-oxidation of FAs in the liver.

In conclusion, this study demonstrates that ADT is effective in improving HFD induced fatty acid metabolism disorders. Specifically, ADT may exert protective effects by: (i) inhibiting overexpression of ACC1, FAS, and SCD1, thereby reducing endogenous synthesis of toxic palmic acid (C16:0) and oleic acid (C18:1); (ii) up-regulating FATPs, L-FABP, CPT1α and Mitofusin1, thus enhancing the β-oxidation of liver fatty acids; (iii) activating FADS1 and FADS2 to improve the desaturated metabolic pathway of EFAs (Figure 4). These results may provide important new insights into the role of ADT in hepatic lipid metabolism, but further researchs are still needed to evaluate its potential application in clinical treatment of hyperlipidemia and NAFLD.

**Figure 4.**
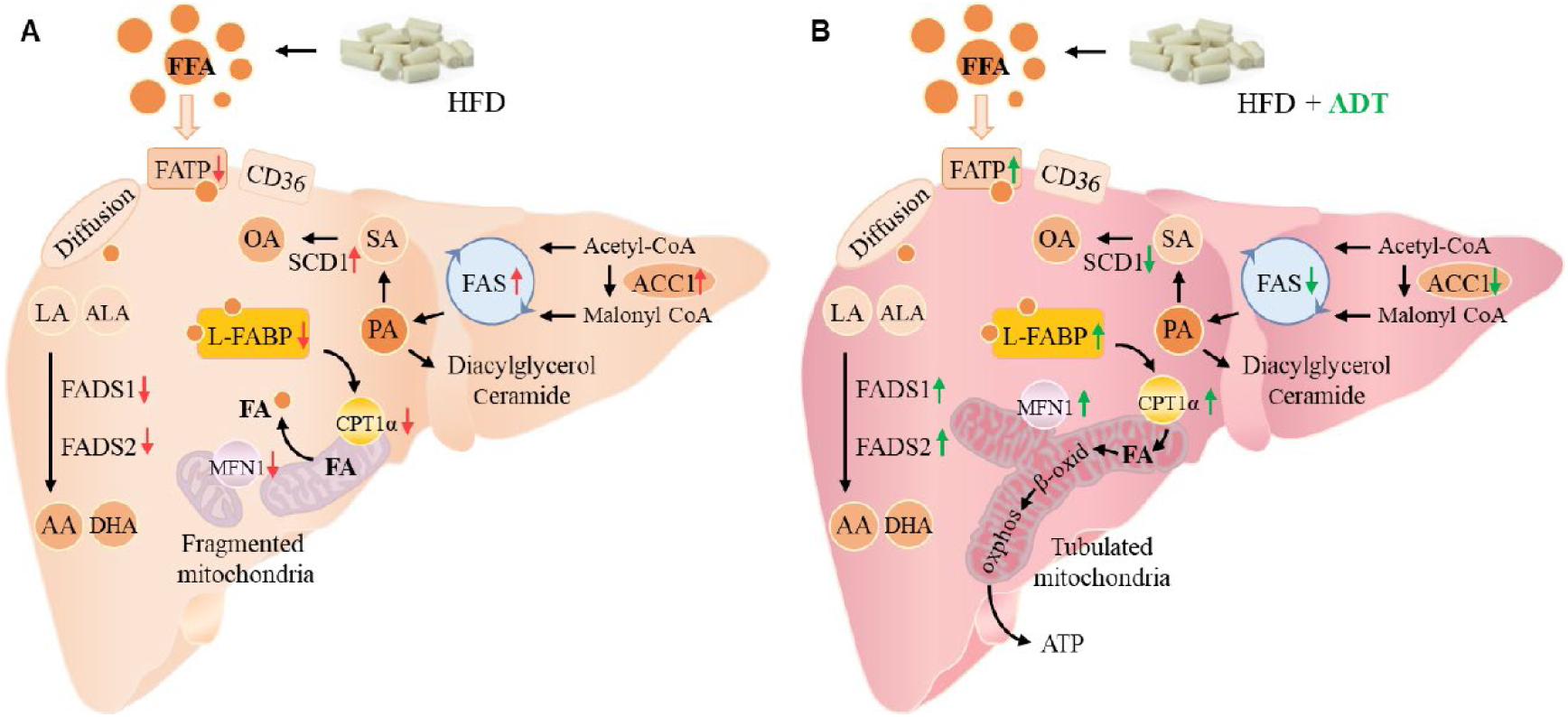
The mechanistic flowchart of ADT on liver FAs metabolism in HFD hamsters. Compared with the HFD group, ADT administration significantly inhibited the overexpression of ACC1, FAS and SCD1, activated FADS1 and FADS2, thereby reducing endogenous synthesis of PA, SA and OA, and improved the desaturated metabolic pathway of EFAs. In addition, ADT upregulated the levels of FATPs, L-FABP, and CPT1α, thus increasing the uptake, binding/isolation, transport, and β-oxidation of FAs. More importantly, ADT obviously increased the expression of Mitofusin1 to promote mitochondrial fusion state and maximal β-oxidation. PA= Palmitic acid; SA=Stearic acid; OA= Oleic acid; EFAs= Essential fatty acids, including LA and ALA; MFN1= Mitofusin1.

## Abbreviations

ADT: anethole dithiolethione, 5-(4-methoxyphenyl)-3H-1,2-dithiole-3-thione
ACC1: acetyl-CoA carboxylase1
CPT1α: carnitine palmitoyltransferase 1α
EFAs: essential fatty acids
FAs: fatty acids
FAS: fatty acid synthase
FADS1: delta-5 desaturases
FADS2: delta-6 desaturases
FAT/CD36: fatty acid transposase
FATP: fatty acid transport protein
H_2_S: hydrogen sulfide
HFD: high-fat diet
L-FABP: liver fatty acid binding protein
Mitofusin1: mitochondrial fusion protein 1
MUFAs: monounsaturated fatty acids
NAFLD: non-alcoholic fatty liver disease
PUFAs: polyunsaturated fatty acids
SFAs: saturated fatty acids
SCD1: stearoyl-CoA desaturase1

## Conflicts Of Interest

The authors declare that they have no competing interests.

